# A novel mouse model of rare neurodevelopmental disorder, TBCK Syndrome

**DOI:** 10.64898/2026.05.07.723566

**Authors:** Ashley J. Melendez-Perez, Emily L. Durham, Dana E. Layo-Carris, Elizabeth M. Gonzalez, Emily E. Lubin, Sarina M. Smith, Kaitlyn E. Worthington, Kaitlin A. Katsura, Rajesh Angireddy, Xiao-Min Wang, Kelly J. Abdalla, Divya Nair, Aaron Black, Abdias Diaz-Rosado, Brianna Ciesielski, W. Timothy O’Brien, Elizabeth J.K. Bhoj

## Abstract

TBCK Syndrome is a rare Mendelian disorder caused by variants in the *TBCK* gene. Although symptoms affect multiple organ systems, hallmark features include intellectual and developmental disability, craniofacial differences, hypotonia, and premature death. At the cellular level, TBCK has been implicated in mTOR signaling, autophagy, mitophagy, and mRNA trafficking; however, the mechanisms underlying disease onset and progression remain unclear. To address this gap, we characterized a mouse model of TBCK Syndrome. These mice lack exon 5 of the *TBCK* gene, resulting in a whole-body knockout of *Tbck*, modeling the most severe known variant. We performed a comprehensive battery of developmental assays, along with microcomputed tomography and histological analyses, which revealed systemic alterations consistent with those observed in affected individuals. Notably, phenotypic changes arising from Tbck loss emerge early and are detectable in the brain, indicating a primary neurodevelopmental origin of disease pathology. Rigorous characterization of this Tbck-deficient mouse establishes the first *in vivo* platform to investigate disease mechanisms and provides a foundation for preclinical evaluation of gene and targeted pharmacological therapy strategies.

**Summary Statement:** This study establishes a rigorously validated animal model recapitulating systemic features of TBCK Syndrome, enabling targeted investigation of disease biology and preclinical assessment of candidate therapies.

## Introduction

TBCK Syndrome (OMIM #616900) is a multisystemic disorder characterized by progressive neurodevelopmental and respiratory issues (Ortiz-Gonzalez et al., 2025). When individuals with biallelic loss-of-function variants in *TBC1 domain-containing Kinase* (*TBCK*) were first reported in 2016, the clinical phenotype was described as an infant-onset constellation of hypotonia with psychomotor delay and “characteristic coarsening of facial features” (Bhoj et al., 2016; Chong et al., 2016; Guerreiro et al., 2016). To date, about 159 affected individuals worldwide are known to have TBCK Syndrome, classifying this an ultra-rare condition (ultra rare < 1 in 50,000 individuals) with a prevalence of less than 1 in 1,000,000 (Baxter et al., 2026; Harari and Humbert, 2020; Ortiz-Gonzalez et al., 2025). As more affected individuals have been identified, TBCK Syndrome has become better understood as a complex condition with a high degree of variation in clinical phenotype that includes severe hypotonia, progressive respiratory and feeding difficulties, seizures, developmental regression and/or neurologic decompensation, spasticity/contractures, frequent fractures, vision changes, and cardiovascular compromise among other clinical features (Beck-Wödl et al., 2018; Bhoj et al., 2016; Chand et al., 2023; Chong et al., 2016; Dai et al., 2022; De Luca-Ramirez et al., 2023; Durham et al., 2023; Guerreiro et al., 2016; Jacob et al., 2024; Liu et al., 2013; Mandel et al., 2017; Mastromoro et al., 2025; Moreira et al., 2021; Ortiz-González et al., 2018; Ortiz-Gonzalez et al., 2025; Sabanathan et al., 2023; Saredi et al., 2020; Tan et al., 2022; Zapata-Aldana et al., 2019).

In addition to the spectrum of clinical features exhibited across the TBCK Syndrome community, a recent analysis of dozens of children with this rare disorder has determined that a genotype-phenotype correlation remains elusive (Dai et al., 2022; Durham et al., 2023; Ortiz-González et al., 2018; Ortiz-Gonzalez et al., 2025; Sabanathan et al., 2023). Twenty-eight unique TBCK Syndrome causing variants have been reported to date (Dai et al., 2022; Durham et al., 2023; Ortiz-González et al., 2018; Ortiz-Gonzalez et al., 2025; Sabanathan et al., 2023). These variants span the gene and protein domains and, while variation in severity of disease has been highlighted in previous publications, true correlations have yet to be found. Further understanding of variant location and disease severity may be better understood through a deeper characterization of TBCK’s overall function and the specific roles of its individual domains. The TBCK protein has been implicated in multiple vital cellular processes including mTOR signaling, autophagy, lysosomal function, mRNA trafficking, and mitochondrial maintenance (Beck-Wödl et al., 2018; Diaz-Rosado et al., 2025; Liu et al., 2013; Moreira et al., 2021; Ortiz-González et al., 2018; Schuhmacher et al., 2023; Tintos-Hernández et al., 2021; Wu and Lu, 2021; Wu et al., 2014). While TBCK contains annotated pseudokinase, TBC (Tre2-Bub2-Cdc16), and rhodanese-like domains based on sequence homology, the functional properties of all three domains remain to be robustly validated through biochemical and structural studies (Cagwin et al., 2025; Durham et al., 2023).

Mechanistic insights into the overall function of the TBCK protein are also crucial to understanding its contribution in the recently described five-subunit endosomal Rab5 and RNA/ribose intermediary (FERRY) complex (Alazami et al., 2015; Hancarova et al., 2019; Loddo et al., 2020; Philips et al., 2017; Rashvand et al., 2022; Rehman et al., 2019; Riffe and Downes, 2025; Schuhmacher et al., 2022; Schuhmacher et al., 2023; Suleiman et al., 2018). The FERRY complex is comprised of TBCK, PPP1R21, FERRY3 (previously C12orf4), CRYZL1, and GATD1. Clinical syndromes have so far been associated with germline variants in three of the five FERRY complex subunits (Riffe and Downes, 2025; Schuhmacher et al., 2022; Schuhmacher et al., 2023). It has been hypothesized that neurogenetic conditions caused by variants in TBCK, PP1R21, and FERRY3 may represent a shared disease class with convergent, therapeutically targetable disease mechanisms; therefore, verifying these linkages will require an orthogonal experimental approach.

Nonetheless, as we continue to advance research into the function of TBCK, individuals living with TBCK Syndrome demonstrate that this is a life-altering and life-limiting condition (Durham et al., 2023; Ortiz-Gonzalez et al., 2025). The current lack of understanding of the pathomechanism of TBCK variants results in affected individuals slowly succumbing to their disease after losing the ability to interact with the outside world and, in many cases, after experiencing recurrent infections and painful recurrent bone fractures (Durham et al., 2023; Ortiz-Gonzalez et al., 2025). Through this work, we take steps towards addressing the current therapeutic gap through the development and interrogation of a mouse model of TBCK Syndrome.

For rare disease research, an accurate validated animal model is a powerful tool. Once characterized, animal models can help understand how the features of a rare syndrome evolve over time, which tissues and organs are affected, and what therapeutically targetable changes occur on a cell and molecular level. Beyond this, animal models allow for the initial evaluation of putative treatment safety and efficacy. Approximately, one third of variants associated with TBCK Syndrome take the form of a nonsense p.R126X variant, a known founder mutation that is prevalent in those of Puerto Rican descent (Ortiz-González et al., 2018; Ortiz-Gonzalez et al., 2025; Tintos-Hernández et al., 2021). Here, we characterize a novel mouse model of TBCK Syndrome generated through the excision of exon 5, which correlates with the location of the p.R126X variant. We hypothesized that, similar to the human phenotype, the effects of a loss of TBCK would be profound, manifest in early in development with a multi-systemic effect, and with a particular impact on neurodevelopment.

## Results

### Molecular characterization and gross phenotypic validation of novel Tbck^-/-^ mouse model

A *Tbck^-/-^* mouse model was generated through the excision of exon 5 to create a frameshift mutation leading to loss of the Tbck mRNA, likely due to nonsense mediated decay **(Figure 1A)**. Exon 5 was excised because it correlates with the location of the TBCK p.R126X founder variant. The resulting *Tbck^+/+^, Tbck^+/-^,* and *Tbck^-/-^* mice were then molecularly characterized. First, RT-qPCR was performed on cDNA synthesized from RNA extracted from brain tissue pooled across male and female mice for comparison between genotypes **(Figure 1B)**. Primers were designed with two objectives. The first two primer sets (exons 1–5 and exons 4–6) were intended to confirm the absence of exon 5 in the knockout mice. Additional primers were placed toward the 3′ end of the gene, in regions shared across most isoforms, to assess whether alternative transcripts might be generated from an internal start site **(Figure 1A, 1B)**. Since exon 5 was excised to generate the model, it was not surprising that there was no amplification of exon 1-5 for *Tbck^-/-^* mice, as well as reduced amplification *Tbck^+/-^* mice compared to *Tbck^+/+^* mice. In all other conditions, amplification of target regions was reduced for *Tbck^-/-^* mice compared to *Tbck^+/-^* mice, which was reduced compared to *Tbck^+/+^* mice. This suggests that an alternative start site could be used to yield other truncated RNAs, which may or may not be translated into protein. Interestingly, quantification of Tbck expression through western blot of protein extracted from lung tissue of *Tbck^+/+^* and *Tbck^-/^*^-^ mice demonstrated the absence of Tbck in *Tbck^-/-^* mice **(Figure 1C)**. This shows that regardless of whether alternative transcription start sites are employed to generate truncated RNA transcripts **(Figure 1B)**, the protein product is not detectable. Thus, molecular phenotyping validates our model at a sub-cellular level.

**Figure 1.**
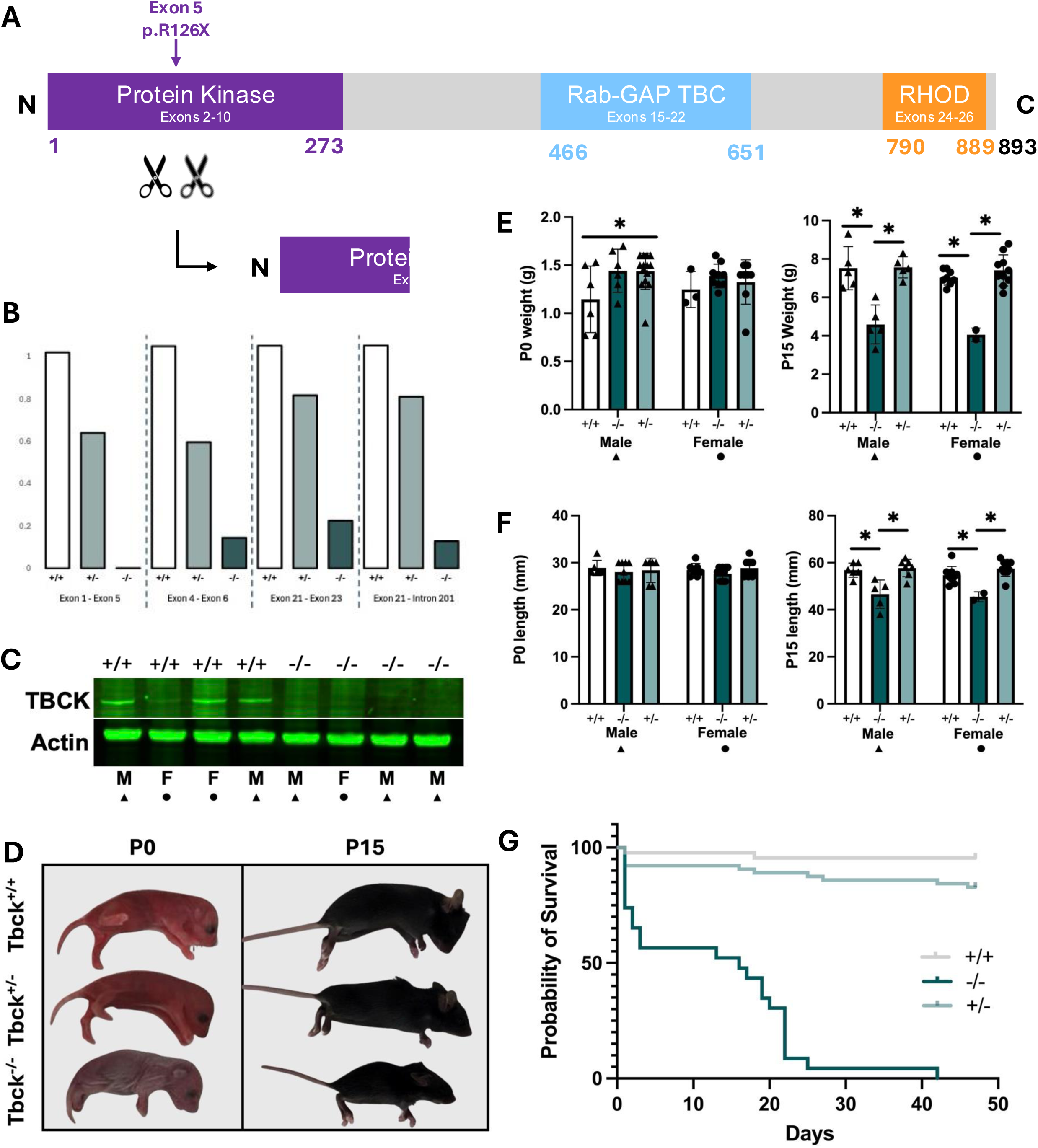
Novel Tbck^-/-^ mouse model shows early gross phenotypic and lifespan differences. **A.** Adapted from Durham et al 2023, schematic of mouse model generation through excision of exon 5, located within the protein kinase domain, to induce frameshift mutation leading to truncated protein. The TBCK founder variant, p.R126X, localizes to exon 5. **B.** RT-qPCR performed on cDNA extracted from brain tissue of Tbck^+/+^ (unfilled bars), Tbck^+/-^ (slate bars), and Tbck^-/-^ (teal bars) mice. Forward and reverse primer combinations were employed to amplify 4 different regions. **C.** Western blot of P0-P1 male and female Tbck^+/+^ and Tbck^-/-^ mice, with Actin used as a loading control. **D.** Pictures of Tbck^+/+^, Tbck^+/-^, and Tbck^-/-^ mice at P0 (left) and P15 (right). **E.** Weights of male and female Tbck^+/+^ (unfilled bars, P0 n= 6M, 3F, P15 n=5M, 8F), Tbck^+/-^ (slate bars, P0 n= 14M, 8F, P15 n=5M, 10F), and Tbck^-/-^ (teal bars, P0 n= 6M, 9F, P15 n=5M, 2F) mice at P0 (left) and P15 (right). * = p<0.05. **F.** Lengths of male and female Tbck^+/+^ (unfilled bars), Tbck^+/-^ (slate bars), and Tbck^-/-^ (teal bars) mice at P0 (left) and P15 (right). * = p<0.05. **G.** Kaplan-Meier survival curve of Tbck^+/+^ (gray), Tbck^+/-^ (slate), and Tbck^-/-^ (teal) mice. n≥7 per genotype

We then performed a general characterization of the mouse model. Recapitulating what was previously demonstrated in both humans and mice (Nair et al., 2023; Wang et al., 2026), *Tbck^+/+^* and *Tbck^+/-^* mice appear similar at both P0 and P15, while *Tbck^-/-^* mice appear to be smaller **(Figure 1D)**. Unexpectedly, there is a statistically significant difference in weight of *Tbck^+/+^* and *Tbck^+/-^* male mice at P0 (p=0.0496) but not between any other genotypes of the same sex (male mice: *Tbck^+/+^* vs *Tbck^-/-^* p=0.1013, *Tbck^+/-^* vs *Tbck^-/-^*p=0.9992; female mice: *Tbck^+/+^* vs *Tbck^-/-^* p=0.4779, *Tbck^+/-^* vs *Tbck^-/-^* p=0.6577, *Tbck^+/+^* vs *Tbck^+/-^* p=0.8432) or between opposite sex mice of the same genotype (male vs female *Tbck^+/+^* p=0.5864; male vs female *Tbck^-/-^* p=0.6081; male vs female *Tbck^+/-^* p=0.1826) **(Figure 1E, left)**. By P15, though, statistically significant differences are noted between *Tbck^-/-^* mice when compared to both *Tbck^+/+^* and *Tbck^+/-^* mice of the same sex (male mice: *Tbck^+/+^* vs *Tbck^-/-^* p=0.0008, *Tbck^+/-^* vs *Tbck^-/-^* p=0.0008, *Tbck^+/+^* vs *Tbck^+/-^* p=0.9977; female mice: *Tbck^+/+^* vs *Tbck^-/-^* p≤0.001, *Tbck^+/-^* vs *Tbck^-/-^* p≤0.001, *Tbck^+/+^* vs *Tbck^+/-^* p=0.4397) **(Figure 1E, right)**. As expected, no sex-based differences were observed between mice of the same genotype (male vs female *Tbck^+/+^* p=0.3871; male vs female *Tbck^-/-^* p=0.3463; male vs female *Tbck^+/-^* p=0.6764). Further, at P0, there was no statistically significant difference in length between any genotypes of the same sex (male mice: *Tbck^+/+^* vs *Tbck^-/-^* p=0.7323, *Tbck^+/-^* vs *Tbck^-/-^* p=0.9506, *Tbck^+/+^* vs *Tbck^+/-^* p=0.9097; female mice: *Tbck^+/+^* vs *Tbck^-/-^*p=0.5238, *Tbck^+/-^* vs *Tbck^-/-^* p=0.1713, *Tbck^+/+^* vs *Tbck^+/-^* p=0.8524) **(Figure 1F, left)**, while, at P15, there were statistically significant differences noted between *Tbck^-/-^* mice when compared to both *Tbck^+/+^* and *Tbck^+/-^* mice of the same sex (male mice: *Tbck^+/+^* vs *Tbck^-/-^*p=0.0302, *Tbck^+/-^* vs *Tbck^-/-^* p≤0.001, *Tbck^+/+^* vs *Tbck^+/-^* p=0.9268; female mice: *Tbck^+/+^* vs *Tbck^-/-^* p=0.009, *Tbck^+/-^* vs *Tbck^-/-^* p≤0.001, *Tbck^+/+^* vs *Tbck^+/-^* p=0.2057) **(Figure 1F, right)**. No differences in length were observed between opposite sex mice of the same genotype at P0 (male vs female *Tbck^+/+^* p=0.6405; male vs female *Tbck^-/-^* p=0.6497; male vs female *Tbck^+/-^*p=0.6897) or P15 (male vs female *Tbck^+/+^* p=0.2377; male vs female *Tbck^-/-^* p=0.736; male vs female *Tbck^+/-^* p=0.9044) **(Figure 1F)**. Finally, Tbck^-/-^ mice displayed reduced survival compared to *Tbck^+/+^* and *Tbck^+/-^* mice **(Figure 1G)**, recapitulating the life-limiting prognosis of TBCK Syndrome. Together, these molecular and gross phenotypic characterizations validate the utility of this mouse model to represent, in a controlled system, the phenotypic signature of TBCK Syndrome.

### Microcomputed Tomography (micro-CT) quantification of craniofacial differences of novel Tbck^-/-^ mouse model

In addition to hypotonia, distinct craniofacial differences are often highlighted in TBCK Syndrome (Bhoj et al., 2016; Durham et al., 2023; Ortiz-González et al., 2018; Ortiz-Gonzalez et al., 2025). Thus, our characterization of this mouse model included an assessment of brain and calvarial form **(Figure 2)**. Gross assessment of embryonic mice indicates that changes in relative size of mice due to loss of Tbck are largely post-natal; however, the lack of Tbck can be observed through changes within the calvarium. Assessment of the ventricles within the brain space using contrast enhanced micro-CT indicates significantly larger ventricular space in Tbck^-/-^ mice as compared to both Tbck^+/+^ (p=0.0014) and Tbck^+/-^ (p=0.0096) littermates while no difference was identified between ventricular volume of Tbck^+/+^ and Tbck^+/-^ littermates (p=0.8446) **(Figure 2B)**. At P0 this increase in the volume of space occupied by the brain held with significantly larger endocast volume being observed in Tbck^-/-^ mice as compared to Tbck^+/+^ (p=0.026) and Tbck^+/-^ (p=0.0021) littermates. Again, there was no difference identified in endocast volume between Tbck^+/+^ and Tbck^+/-^ (p=0.91) mice. In adult mice this held true though there are subtle differences in skull form **(Figure 2)**.

**Figure 2.**
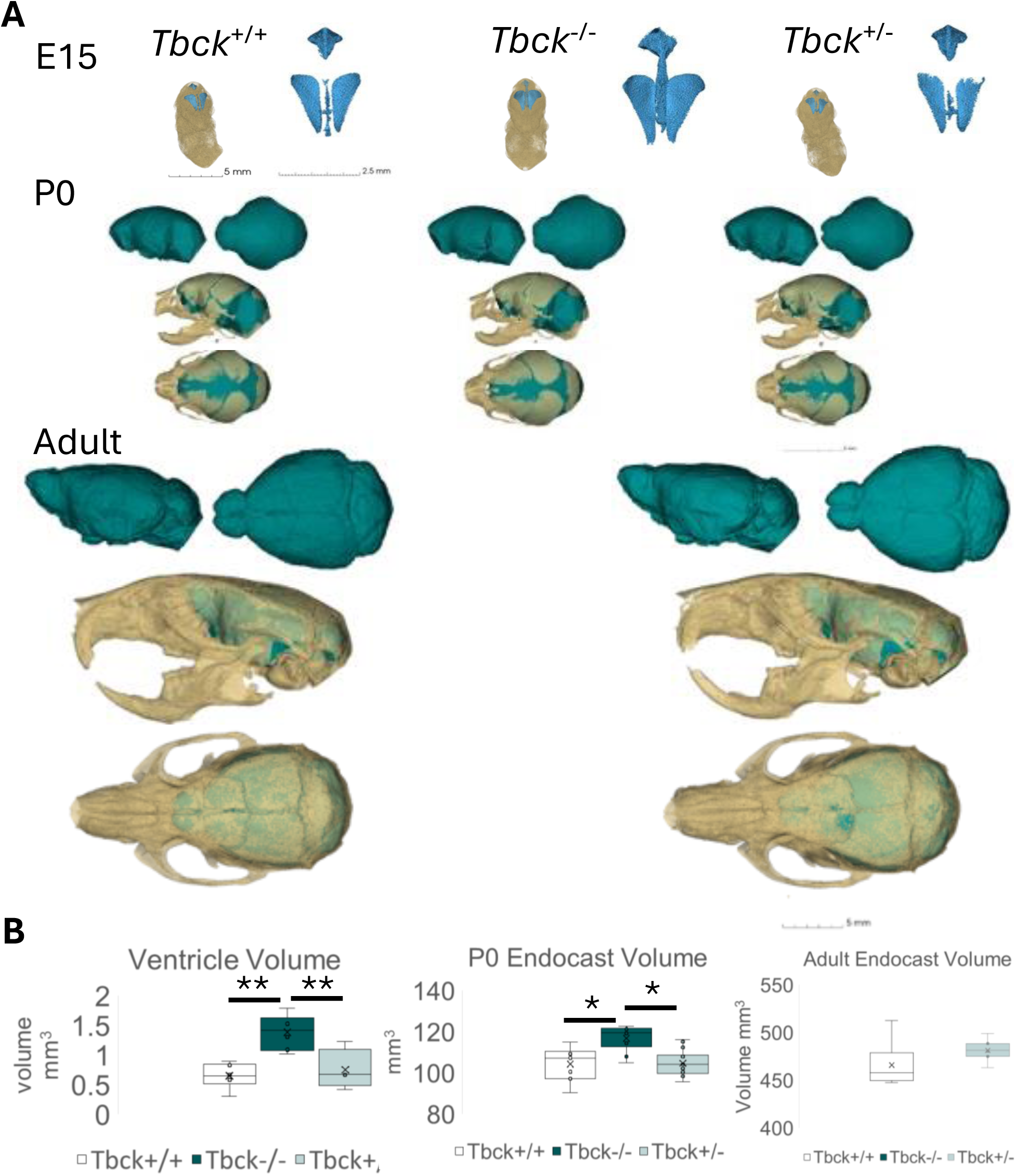
Micro-CT-based analyses of Tbck^-/-^ mice show changes within the calvarium. **A.** Representative 3D Micro-CT renderings of E15 embryos and ventricles (E15, Top Row: Tbck^+/+^ left, Tbck^-/-^ middle, Tbck^+/-^ right) highlights increase ventricular volume in Tbck^-/-^ embryos. Similarly, endocasts (Left lateral and superior views in teal) displaying the space occupied by the brain in P0 mice (P0, Middle Row: Tbck^+/+^ left, Tbck^-/-^ middle, Tbck^+/-^ right) indicate continued if more subtle volumetric changes due to genotype. Only Tbck^+/+^ (Bottom row left), and Tbck^+/-^ (Bottom Row right) mice survive to adulthood and assessment of endocasts (left lateral and superior views in teal) indicate no significant effect genotype on craniofacial form. Skulls with endocast highlighted within displayed for each genotype with left lateral view above superior view. **B.** Quantification of ventricle (left), P0 endocast (middle), and adult endocast (right) volume. n≥4 per genotype per assessment. *p≤0.05, ** p≤0.01 Scales indicated for each age on figure.

### Histological assessment of neurodevelopmental targets and brain development of the TBCK Syndrome mouse model

Brain structure anomalies and neurodevelopmental symptoms are almost ubiquitous in individuals affected with TBCK Syndrome (Durham et al., 2023; Ortiz-Gonzalez et al., 2025). Thus, we extended our analysis of Tbck^-/-^ mice to an in-depth assessment of the brain. In P0 mice, no structural differences were noted between genotypes using basic Hematoxylin & Eosin (H&E) staining for morphology or Cresyl Violet (CV) staining for brain cytoarchitecture **(Figure 3A)**. Analysis for neuronal targets (SOX2, NeuN) determined no difference between genotypes (SOX2 p=0.105, NeuN p=0.693) (**Figure 3B-C**). Using MAP2 and GFAP, we assessed presence of neuronal and glial cells within a standard region of interest and identified significant differences in MAP2 (p=0.0254) between Tbck^+/+^ and Tbck^+/-^ mice (p=0.0193) (**Figure 3B-C**). We also observed a significant increase in the number of GFAP positive cells in Tbck^+/-^ mice as compared to Tbck^+/+^ mice (p=0.0193) and Tbck^-/-^ mice as compared to Tbck^+/+^ mice (p=0.0488) (**Figure 3B-C**). No significant differences between genotypes in Iba1 staining were identified at this age **(Figure 3C)**.

**Figure 3.**
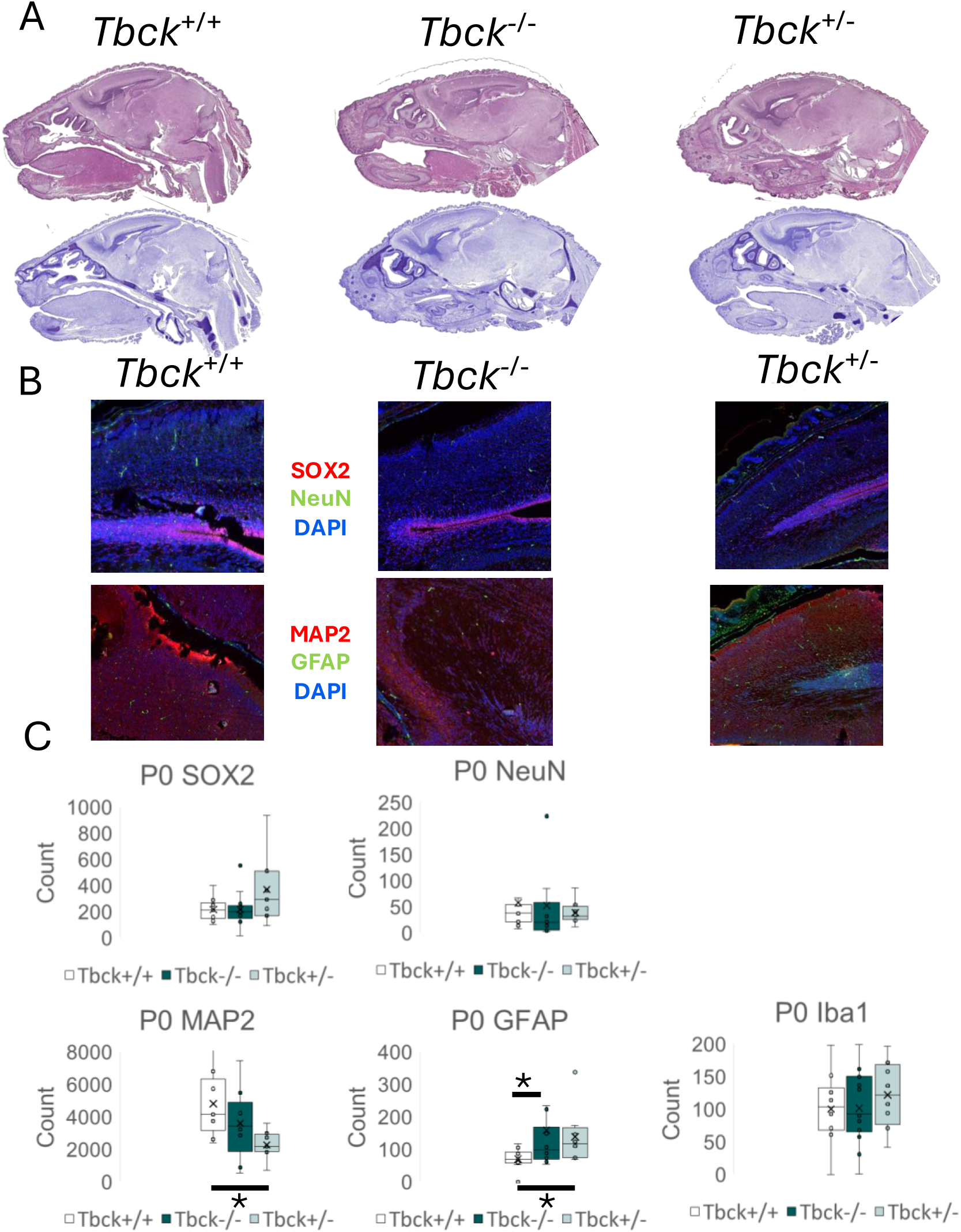
Histological assessment of P0 mice indicates modulation to within-brain targets important to neurodevelopment. **A.** Representative approximately midline sections of Tbck^+/+^ (left), Tbck^-/-^ (middle), Tbck^+/-^ (right) mice indicate no significant morphological changes by genotype in either basic H&E stain (top) or brain specific Nissl staining (bottom). B-C. **B.** Using a standard region of interest displayed, count of stained cells for SOX2 (Red) and NeuN (Green) (top row) indicates no differences in these targets by genotype Tbck^+/+^ (left), Tbck^-/-^ (middle), Tbck^+/-^ (right). Decreased signals for MAP2 (red) and increased signals for GFAP (green) for diminished TBCK confirm changes to neurodevelopmental targets by genotype. **C.** Quantification of target signal by count of cells within a standard region of interest indicates decreased MAP2 (Red) in Tbck^+/-^ mice compared to Tbck^+/+^ mice. For GFAP (green) increased signal in Tbck^-/-^ and Tbck^+/-^ mice compared to control Tbck^+/+^ mice highlights changes due to levels of TBCK expression in the brain during development. No significant differences between genotypes were identified in microglia via Iba1 staining. n≥4 per genotype per assessment *p≤0.05.

Knowing that brain growth and development occur at an exponential rate postnatally (Richtsmeier and Flaherty, 2013), we assessed these targets in P15 mice as well. Morphological (H&E) and cytoarchitectural (CV) staining reveal no grossly significant differences between genotypes **(Figure 4A)**. In correlation with these results, we did not find significant differences in neuronal targets (p=0.4834 SOX2, p=0.7061 NeuN) (**Figure 4B-C**) or cerebellar targets (p=0.7239, Calbindin) between genotypes (**Figure 4C**). However, we did note high degrees of variability aligning with active neurodevelopment. Assessment of MAP2 and GFAP identified modulation of these neuronal targets with presence/absence of Tbck **(Figure 4B)**. Within the region of interest assessed, MAP2 (p=0.047) was increased in Tbck^-/-^ mice as compared to Tbck^+/+^ (p=0.004) and Tbck^+/-^ (p=0.003) mice **(Figure 4B-C)**. For GFAP (p=0.002), significant differences were noted between Tbck^+/+^ and Tbck^+/-^ mice (p=0.035) and Tbck^-/-^ and Tbck^+/-^ mice (p=0.001) but not between Tbck^+/+^ and Tbck^-/-^ mice (p=0.5805) (**Figure 4B-C**). No significant differences between genotypes were identified for the microglia marker Iba1 (**Figure 4C**).

**Figure 4.**
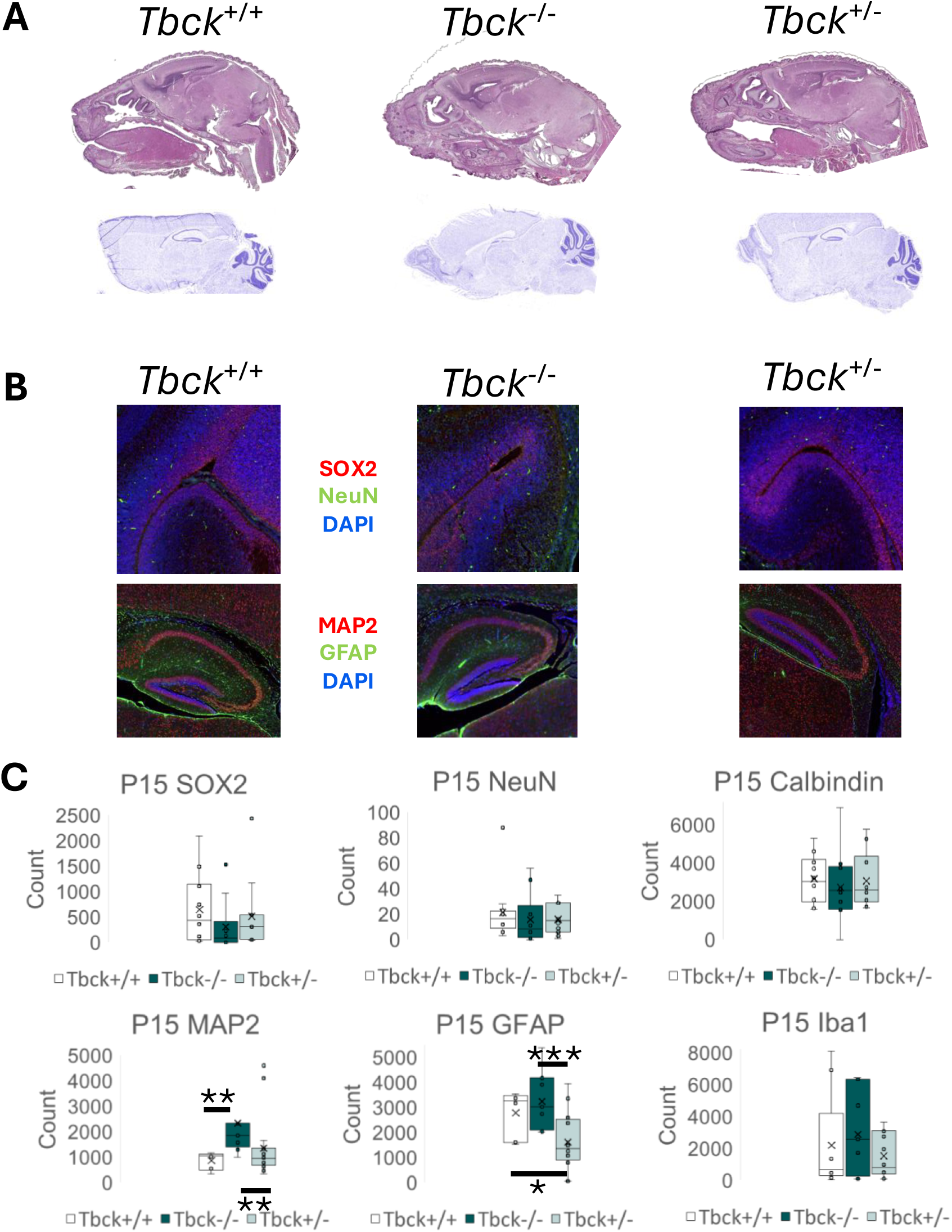
Histological assessment of P15 mice indicates significant changes to within-brain targets important to neurodevelopment. **A.** Representative approximately midline sections of Tbck^+/+^ (left), Tbck^-/-^ (middle), Tbck^+/-^ (right) mice indicate no significant morphological changes by genotype in either basic H&E stain (top) or brain specific Nissl staining (bottom). **B.** Using a standard region of interest displayed, count of stained cells for SOX2 (Red) and NeuN (Green) (top row) indicates no differences in these targets by genotype Tbck^+/+^ (left), Tbck^-/-^ (middle), Tbck^+/-^ (right). Increased signals for MAP2 (red) and GFAP (green) for Tbck^-/-^ (middle) mice (bottom row) confirm changes to neurodevelopmental targets by genotype. **C.** Quantification of target signal by count of cells within a standard region of interest indicates increased MAP2 (Red) in Tbck^-/-^ mice compared to Tbck^+/+^ and Tbck^+/-^ mice. For GFAP (green) reduced signal in Tbck^+/-^ mice compared to both other genotypes highlights changes due to levels of TBCK expression in the brain during development. No significant differences between genotypes were identified for Calbindin or Iba1. n≥4 per genotype per assessment *p≤0.05 ** p≤0.01 ***p ≤0.001.

### Transcriptomic profiling of Tbck^+/+^ versus Tbck^-/-^ mice

To broadly interrogate the resulting transcript-level dysregulation in *Tbck^-/-^* mice, we performed bulk RNA-sequencing that compared brain samples from P0-P1 *Tbck^+/+^* and *Tbck^-/-^* mice. Reassuringly, *Tbck* was the differentially expressed gene (DEG) with the greatest log2fold change **(Figure 5A)**. A total of 13 genes were significantly differentially expressed (2 downregulated, 11 upregulated) **(Figure 5A, subpanel;** **Table 1)**. Notably, *Tbck* was the only FERRY complex gene to show significant differential expression between the two conditions **(Figure 5B)**. Further, there was not clear delineation between the *Tbck^+/+^* and *Tbck^-/-^* mice based on principal component analysis (PCA) when looking at the top 500 DEGs **(Figure 5C, left)**, which was surprising based on the distinct features at the levels of gross phenotyping **(Figure 1D-G)**, micro-CT **(Figure 2)**, and histologically **(Figures 3-4)**. Clear groupings emerged when the PCA was restricted to the top 50 DEGs **(Figure 5C, right)**, indicating that this subset of genes may be driving the profound phenotypic differences.

**Figure 5.**
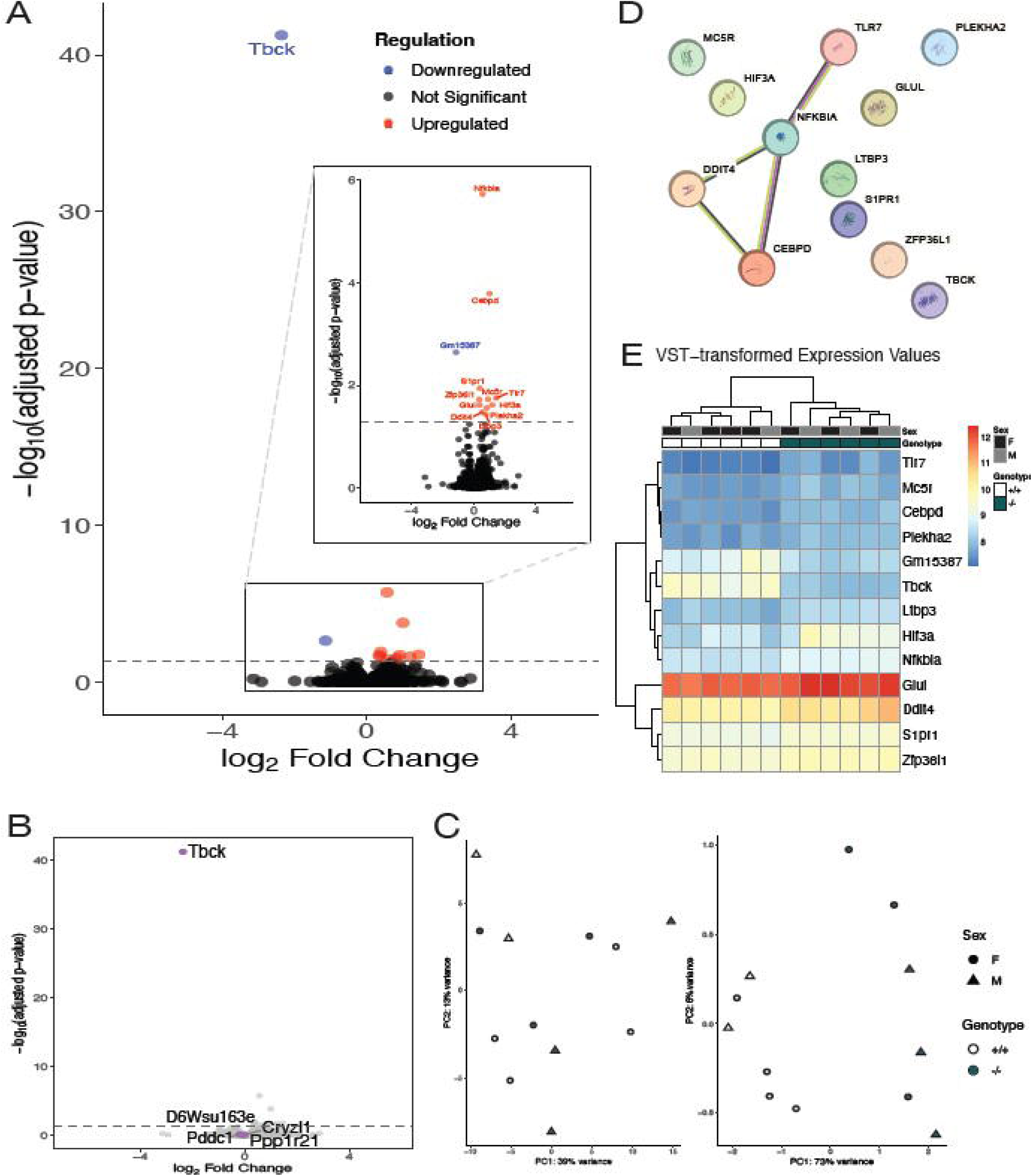
Transcriptomic profile of Tbck^-/-^ mice. **A.** Volcano plot demonstrating differentially expressed genes (DEGs) (red = upregulated, blue = downregulated, black = no significant change) between Tbck^+/+^ and Tbck^-/-^ mice. Subpanel magnifies DEGs within the black box. **B.** Volcano plot highlighting FERRY Complex genes in magenta. **C.** Principal component analyses of the top 500 DEGs (left) and top 50 DEGs (right). Circle = female mice, triangle = male mice, unfilled shape = Tbck^+/+^ mice, teal shape = Tbck^-/-^ mice. **D.** STRING database protein-protein association network analysis and functional enrichment analyses of DEGs. **E.** Heatmap of DEGs. Black = female mice, gray = male mice, unfilled = Tbck^+/+^ mice, teal = Tbck^-/-^ mice.

**Table 1.**
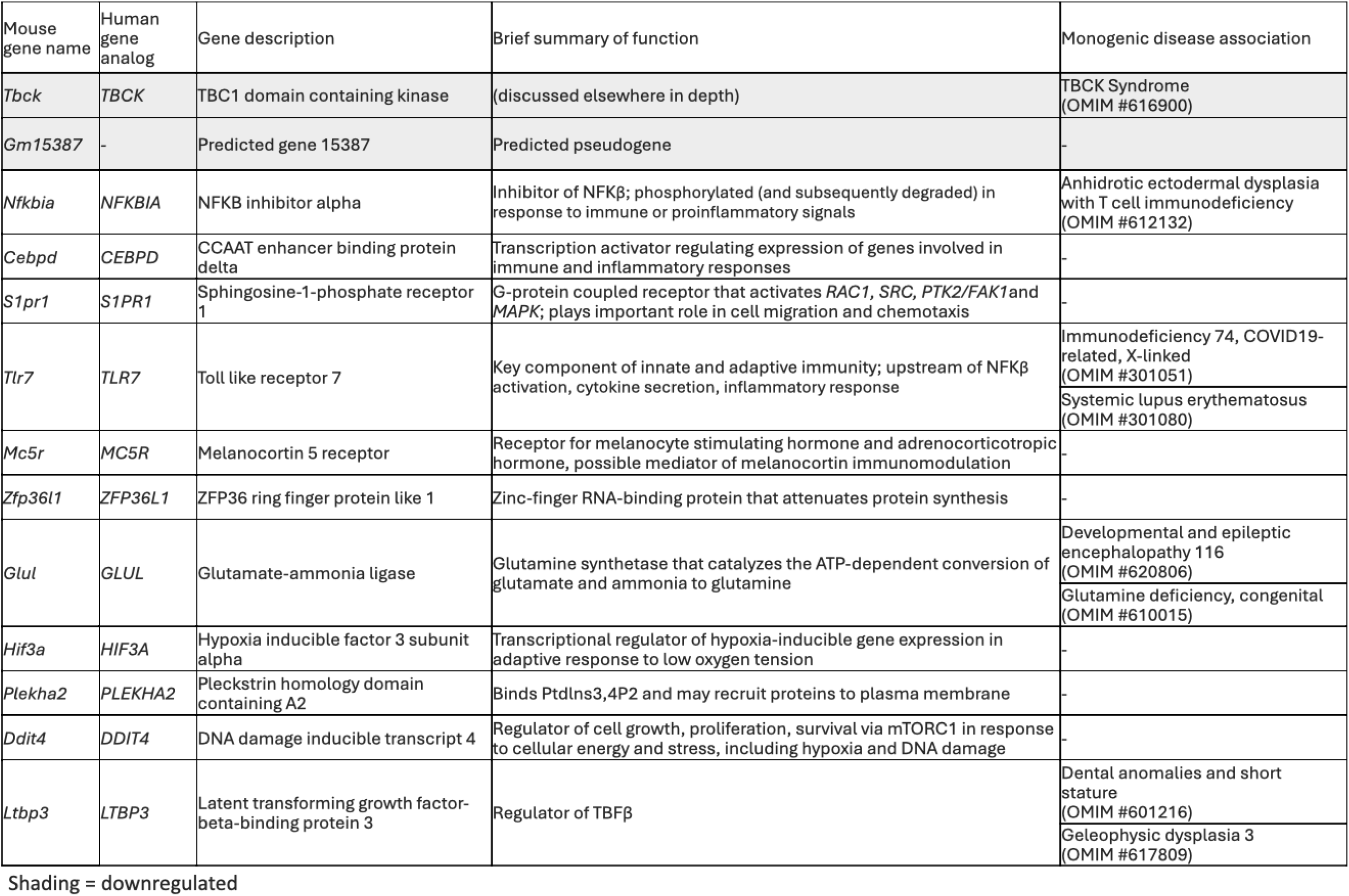
Overview of differentially expressed genes

To better understand the relationship between the DEGs that reached significance, we performed a STRING database protein functional enrichment analysis to identify putative disease-relevant protein interaction networks **(Figure 5D)**. *Gm15387,* a predicted pseudogene differentially expressed between the two conditions, was not able to be included in this protein-level analysis as it lacks a protein correlate **(Table 1)**. *Nfkbia, Tlr7, Ddit4,* and *Cebpd,* genes whose expression is regulated by stress signals like pro-inflammatory or hypoxic cues, form one cluster **(Figure 5D**, **Table 1)**. Interestingly, *Hif3a* is another gene implicated in hypoxic response. Prior work in neurons, astrocytes, and microglia from primary mouse brain cultures have shown that the expression of *Ddit4* and *Hif3a* are sensitive to conditions of decreasing oxygen content (Ravel-Godreuil et al., 2024). So, even though we were unable to delineate our RNA-sequencing data at a cell type-specific resolution, we generated a heatmap of genes to understand the potential clustering on the basis of hypoxia sensitivity **(Figure 5E)**.

## Discussion

TBCK Syndrome is a uniformly fatal, life-limiting pediatric neurodegenerative condition. Community partnerships through The TBCK Foundation have led to patient registries, natural history studies, and family meetings, which have facilitated information-and resource-sharing, along with crucial advocacy efforts. These heroic grass-roots initiatives have parried some of the most significant barriers around progress in pediatric neurodegenerative disorders, namely a lack of both research attention and public awareness (Baxter et al., 2026; Elvidge et al., 2025). Nonetheless, there remains a gap between our ability to diagnose or longitudinally phenotype individuals with TBCK Syndrome and our ability to offer therapeutic interventions beyond symptom management and palliative interventions.

Since the initial description of biallelic loss-of-function variants in *TBCK*, increasing numbers of affected individuals have been identified, revealing a multisystem disorder with variable severity but consistently severe neurological involvement. However, despite improved clinical characterization, the biological mechanisms underlying TBCK deficiency remain poorly understood and no disease-modifying therapies currently exist. Modeling of TBCK Syndrome in mice has been challenging. A subtle phenotype was identified in Tbck^+/-^mice characterized by reduced exploratory and rearing behavior (Nair et al., 2023). Complete knockout of TBCK in a different model caused perinatal lethality (Wang et al., 2026). In this study, we present a novel *Tbck* knockout mouse model that recapitulates key clinical and anatomical features of TBCK Syndrome, establishing an experimentally tractable system for investigating disease biology and evaluating candidate therapeutic strategies.

A key contribution of this work is the development and validation of a genetically relevant model of TBCK loss-of-function. The engineered excision of exon 5 produces a frameshift leading to nonsense-mediated decay of *Tbck*, functionally mimicking the recurrent p.R126X founder variant that accounts for a substantial proportion of affected individuals worldwide. Molecular analyses confirmed the absence of Tbck protein in homozygous animals, while phenotypic characterization revealed reduced body size and markedly decreased survival relative to littermate controls **(Figure 1C-G)**. These findings align with prior work demonstrating that TBCK plays an important role in regulating cellular growth and proliferation (Liu et al., 2013; Nair et al., 2023; Ortiz-González et al., 2018; Tintos-Hernández et al., 2021; Wu and Lu, 2021). *In vitro* depletion of TBCK leads to reduced cell size, impaired proliferation, and disruption of cytoskeletal organization through suppression of the mechanistic target of rapamycin (mTOR) signaling pathway (Liu et al., 2013; Wu and Lu, 2021; Wu et al., 2014). Because mTOR signaling is a central regulator of cellular growth and metabolic homeostasis, dysregulation of this pathway may contribute to the growth deficits and developmental abnormalities observed both in this model and in affected individuals.

The neuroanatomical abnormalities observed in the *Tbck*-deficient mice further support the translational relevance of this model. Micro-CT revealed enlarged ventricular spaces and altered craniofacial morphology, findings that parallel neuroimaging observations in individuals with TBCK Syndrome **(Figure 2)**. Early clinical reports describing TBCK deficiency documented hypotonia, intellectual disability, and characteristic neuroimaging features including brain atrophy and white-matter abnormalities (Bhoj et al., 2016; Chong et al., 2016; Ortiz-González et al., 2018). Structural brain abnormalities observed in the present mouse model therefore reinforce the concept that TBCK plays an important role in normal neurodevelopment and brain structural integrity.

Growing evidence suggests that TBCK functions at the intersection of several cellular pathways critical for neuronal homeostasis, including mTOR signaling, autophagy, lysosomal function, and intracellular trafficking (Beck-Wödl et al., 2018; Diaz-Rosado et al., 2025; Liu et al., 2013; Moreira et al., 2021; Ortiz-González et al., 2018; Schuhmacher et al., 2023; Tintos-Hernández et al., 2021; Wu and Lu, 2021; Wu et al., 2014). Early studies demonstrated that TBCK regulates expression of components of the mTOR complex, with knockdown leading to reduced mTOR signaling activity and impaired cell growth (Liu et al., 2013). Subsequent studies in patient-derived fibroblasts revealed abnormalities in autophagy and mitochondrial quality control, including accumulation of mitophagosomes, reduced mitochondrial respiratory capacity, and decreased mitochondrial DNA content (Tintos-Hernández et al., 2021). These defects appear to arise secondary to impaired lysosomal proteolytic function, linking TBCK deficiency to disruption of the lysosome–mitochondria axis. Such findings are consistent with neuropathological observations suggesting that TBCK-associated disease may share mechanistic features with lysosomal storage disorders and other neurodegenerative conditions characterized by impaired autophagy (Ortiz-González et al., 2018).

The neuropathological assessments of Tbck^-/-^ mice at P0 and P15 reveal a temporally dynamic pattern of neuronal and glial perturbation that aligns with the known clinical burden of brain abnormalities in TBCK syndrome (Durham et al., 2023; Ortiz-Gonzalez et al., 2025). At P0, despite the absence of gross morphological or cytoarchitectural differences that are consistent with the early-onset but structurally subtle nature of TBCK-related neuropathology, a significant reduction in MAP2 signaling was observed across in Tbck^+/-^ as compared to Tbck^+/+^ (**Figure 3C**). MAP2 is the predominant cytoskeletal regulator within neuronal dendrites, and its expression is closely tied to dendritic arborization and synaptic function (DeGiosio et al., 2022), suggesting that even partial TBCK haploinsufficiency may subtly impair early dendritic maturation. By P15, a period coinciding with the murine brain growth spurt during which rapid expansion of dendritic arbors from newly differentiated neurons occurs to accommodate emerging synaptic connections, peaking between approximately P10-12 (Laeremans et al., 2013), the pattern shifted whereas by P15 Tbck^-/-^ mice exhibited a significant increase in MAP2 signal compared to Tbck^+/+^. In contrast, Tbck^+/-^ mice no longer differed from Tbck^+/+^. This paradoxical upregulation may reflect a compensatory or dysregulated dendrite elaboration response in the context of TBCK-associated lysosomal and mTOR pathway dysfunction (Flores-Mendez et al., 2025; Tintos-Hernández et al., 2021). Concurrent with the MAP2 changes at P15, GFAP, a well-established marker whose upregulation is characteristic of reactive astrogliosis in response to neuronal stress and pathological CNS conditions (Jurga et al., 2021), was significantly modulated in a genotype-dependent manner, with Tbck^+/+^ mice displaying distinctly different GFAP levels from both heterozygous and homozygous knockout animals. This intermediate and divergent glial phenotype in heterozygotes at P15 is notable and consistent with prior evidence that haploinsufficiency at the TBCK locus can produce a measurable neurologic phenotype (Flores-Mendez et al., 2025; Tintos-Hernández et al., 2021). The absence of significant Iba1 differences at either timepoint suggests that microglial activation is not a predominant feature of early Tbck-loss neuropathology in this model, distinguishing the glial response seen here from classical neuroinflammatory processes. Taken together, these data provide the first *in vivo* evidence of early-onset, progressive neuronal and astroglial perturbation in Tbck^-/-^ mice, establishing a tractable model for interrogating the neurodevelopmental trajectory of TBCK syndrome.

Recent work has further implicated TBCK in neuronal endolysosomal trafficking and mRNA transport (Flores-Mendez et al., 2025; Schuhmacher et al., 2022; Schuhmacher et al., 2023; Wang et al., 2026). In human neuronal models derived from induced pluripotent stem cells, TBCK localizes to endolysosomal vesicles and participates in the FERRY complex, a multiprotein assembly that mediates mRNA transport along endosomal compartments. Loss of TBCK disrupts lysosomal trafficking and leads to reduced axonal mRNA localization, processes essential for maintaining neuronal structure and synaptic function (Flores-Mendez et al., 2025). These findings suggest that TBCK deficiency may contribute to neurodegeneration through impaired intracellular trafficking and disruption of localized protein synthesis in neuronal processes. The severe neurologic phenotype observed in the *Tbck^-/-^* mice is therefore consistent with disruption of fundamental neuronal maintenance pathways.

Transcriptomic analysis of neonatal mouse brain tissue identified a relatively small number of differentially expressed genes between *Tbck*^+/+^ and *Tbck*^-/-^ animals **(Figure 5A)**. While initially surprising given the robust phenotypic differences observed at the organismal and anatomical levels, several factors may explain this finding. The relatively small number of differentially expressed genes in RNA-seq from P0 mouse brains may reflect both the high natural variability in gene expression during the perinatal period and the heterogeneity in phenotypic severity **(Figure 1G)**, such that harvested P0 brains may represent animals at very different stages along a trajectory toward lethality, from hours to weeks before perinatal demise (Kang et al., 2011). Moreover, bulk RNA-sequencing averages transcriptional signals across heterogeneous populations of neurons and glia, potentially obscuring cell type-specific transcriptional responses to TBCK loss. Indeed, recent studies of neuronal TBCK deficiency have highlighted cell type-specific effects on autophagy and mTOR signaling that may not be fully captured by bulk transcriptomic approaches (Flores-Mendez et al., 2025).

Nevertheless, functional network analysis of the differentially expressed genes identified clusters associated with stress and inflammatory signaling pathways, including genes such as *Nfkbia*, *Tlr7*, and *Ddit4* **(Figure 5D)**. These genes are well known mediators of cellular responses to metabolic and environmental stress. Notably, DDIT4 (REDD1) is a stress-induced inhibitor of mTOR signaling, providing a potential mechanistic link between stress responses and metabolic dysregulation in TBCK deficiency **(Table 1)**. The enrichment of genes associated with hypoxia and inflammatory signaling pathways therefore suggests that loss of TBCK may trigger a coordinated cellular stress response during early brain development. Such responses may reflect secondary consequences of mitochondrial dysfunction, lysosomal impairment, or metabolic stress arising from disrupted mTOR signaling.

Importantly, the phenotypic features observed in the *Tbck*^-/-^ mouse parallel aspects of other pediatric neurodegenerative disorders characterized by defects in lysosomal and autophagic pathways. Lysosomal storage disorders and related conditions frequently present with neurodevelopmental impairment, progressive neurologic decline, and structural brain abnormalities (Boustany, 2013; Platt et al., 2018). In TBCK-deficient cells, lysosomal dysfunction leads to accumulation of lipid droplets and impaired degradation of autophagosomes, supporting the concept that defective lysosomal degradation represents a central mechanism underlying disease pathogenesis (Flores-Mendez et al., 2025). The *Tbck*-deficient mouse therefore provides a valuable *in vivo* model to further investigate how disruption of lysosomal and mitochondrial homeostasis contributes to neurodegeneration.

Beyond mechanistic insights, the model described here provides an important platform for translational research. At present, clinical management of TBCK Syndrome is limited to supportive care, reflecting the broader therapeutic gap faced by many ultra-rare pediatric neurodegenerative conditions. Preclinical models are essential for evaluating potential therapeutic approaches and for understanding the natural history of disease progression. Notably, prior work has demonstrated that activation of the mTOR pathway in TBCK-deficient cells remains responsive to amino acid stimulation, including leucine supplementation, suggesting that targeted metabolic interventions may represent a potential therapeutic avenue (Bhoj et al., 2016; Diaz-Rosado et al., 2025; Nair et al., 2023). The availability of a validated mouse model therefore creates opportunities to evaluate candidate metabolic or molecular therapies *in vivo*.

This study has several limitations. Transcriptomic analyses were performed using bulk RNA-sequencing, which limits the ability to detect cell type-specific molecular changes (Hwang et al., 2018). Single-cell transcriptomic or spatial transcriptomic approaches may provide improved resolution of disease-relevant pathways in specific neuronal and glial populations. Additionally, the current analyses focused primarily on early developmental timepoints, whereas TBCK Syndrome in humans often follows a progressive neurodegenerative course. Longitudinal studies across developmental stages will therefore be important to understand how TBCK deficiency drives disease progression over time. Finally, although this model recapitulates several key features of the human disorder, comparison with other emerging TBCK animal models modeling additional disease-causing variants will be necessary to determine the full spectrum of phenotypic variability associated with TBCK loss (Nair et al., 2023; Wang et al., 2026). Importantly, when the variant was crossed into the C57B/6J background, we found that complete deletion of Tbck was lethal (Nair et al., 2023). Thus, mice used in this study were in the C57B/6N background.

In summary, we report the characterization of a novel *Tbck* knockout mouse that recapitulates key phenotypic features of TBCK Syndrome. By providing an experimentally tractable system for investigating TBCK biology *in vivo*, this model offers a critical tool for elucidating the molecular mechanisms underlying this ultra-rare neurodevelopmental disorder. Future studies leveraging this model will enable deeper investigation of TBCK-dependent pathways in neuronal development and may facilitate the development of targeted therapeutic strategies for affected individuals.

## Materials and Methods

### Mouse colony management

*Tbck^tm1a(EUCOMM)Hmgu^* mice on a C57BL/6N genetic background were generated through the Baylor College of Medicine Knockout Mouse Phenotype Project and made available through the International Mouse Phenotyping Consortium. It is characterized as a first allele knock out (reporter-tagged insertion with conditional potential) line. The Jackson Laboratory Phenotyping Center has reported preweaning lethality in homozygote mice.

### Sex and allele genotyping

Mouse genomic DNA was extracted using the HotSHOT method. Briefly, the alkaline lysis buffer contained 25 mM NaOH and 0.2 mM disodium EDTA (pH 12), and the neutralizing buffer consisted of 40 mM Tris-HCl (pH 5). Tissue samples (2-mm ear punches or ∼0.2-cm tail tips) were collected into PCR strip tubes and incubated in 60 µL of alkaline lysis buffer at 95 °C for 30 minutes. Samples were then cooled to 4 °C, and an equal volume of neutralizing buffer was added. The lysates were centrifuged at maximum speed for 2–5 minutes to pellet debris. DNA concentration in the supernatant was measured using a NanoDrop 8000 UV–Vis spectrophotometer.

PCR amplification using sex-specific primers (FWD: GATGATTTGAGTGGAAATGTGAGGTA and REV: CTTATGTTTATAGGCATGCACCATGTA) was performed using KAPA2G Fast HotStart Genotyping Mix reagent (catalog #KK5609) in a total reaction volume of 20 µL. Each reaction contained 10 µL of KAPA master mix, 1 µL of forward primer, 1 µL of reverse primer, 2 µL of template cDNA, and 6 µL of nuclease-free water. Thermal cycling was carried out under the following conditions: an initial denaturation at 94°C for 3 minutes, followed by 33 cycles of denaturation at 94°C for 30 seconds, annealing at 60°C for 30 seconds, and extension at 72°C for 1 minute. A final extension step was performed at 72°C for 10 minutes. PCR products were separated on a 2% agarose gel at 120V for 20 minutes and visualized using a ChemiDoc imaging system (Bio-Rad). Female samples exhibited a single band at 686 bp, whereas male samples showed two bands corresponding to the X- and Y-chromosome products (686 bp and 475 bp) respectively.

Similarly for allele genotyping, the PCR amplification using TBCK locus specific primers (TBCK_WT_FWD: TGTTTGCAGCCATGAGAATC, TBCK_WT_REV: TGCCACAGGATACCCAAAGA, TBCK_KO_FWD: CGGTCGCTACCATTACCAGT, TBCK_KO_REV: CCTCCAATGTTCACAATGAGTT) was performed using KAPA2G Fast HotStart Genotyping Mix reagent (catalog #KK5609) in a total reaction volume of 10 µL. Each reaction contained 5 µL of KAPA master mix, 0.5 µL of each of the four primers above mentioned, 1 µL of template cDNA, and 2 µL of nuclease-free water. Thermal cycling was carried out under the following conditions: an initial denaturation at 95°C for 3 minutes, followed by 10 cycles of denaturation at 95°C for 10 seconds, annealing at 65°C for 15 seconds, and extension at 72°C for 15 seconds, followed by 20 cycles of denaturation at 95°C for 10 seconds, annealing at 60°C for 15 seconds, and extension at 72°C for 15 second. A final extension step was performed at 72°C for 10 minutes. PCR products were separated on a 1.5% agarose gel at approximately 140V and visualized using a ChemiDoc imaging system (Bio-Rad). Wildtype (WT) samples exhibited a single band at 400 bp, whereas homozygous knockouts samples showed a single band at 580bp, and finally, heterozygous samples displayed both bands at 400bp and 580bp.

### RT-qPCR

Total RNA was isolated from whole brain tissue of P0–P1 mice using the Maxwell® RSC SimplyRNA Tissue Kit (AS1340; Promega) in accordance with the manufacturer’s instructions. RNA concentration and purity were assessed using a NanoDrop 8000 UV–Vis spectrophotometer (Thermo Fisher Scientific). Complementary DNA (cDNA) was synthesized from 1 µg of total RNA using the SuperScript™ VILO™ cDNA Synthesis Kit (Thermo Fisher Scientific, cat. 11754050). The resulting cDNA (20 µL reaction) was diluted with 80 µL nuclease-free water prior to downstream applications.

Quantitative PCR was performed using 2X PowerUp™ SYBR™ Green Master Mix (Applied Biosystems, cat. A25741) on an Applied Biosystems QuantStudio™ 3 system. Each reaction contained 2 µL of diluted cDNA, 1 µM primers, and 1X master mix in a total volume of 15 µL. Gene expression levels were normalized to GAPDH, and relative expression was calculated using the comparative Ct (ΔΔCt) method. All samples were run in technical replicates, and four biological replicates were included for each genotype. Primer sequences are provided in **Table 2**.

**Table 2:**
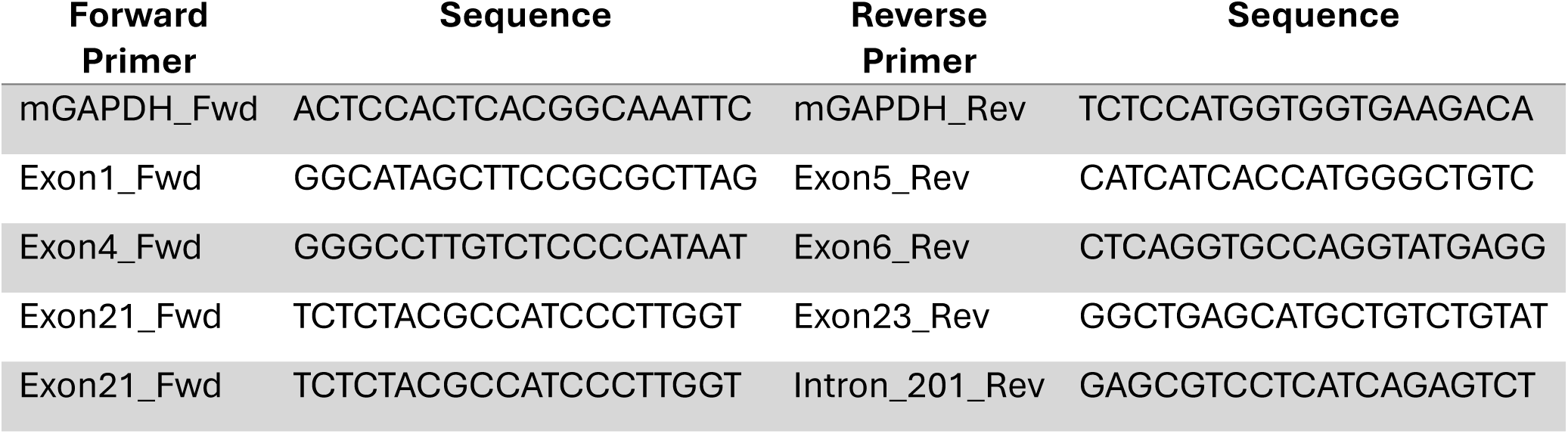
Primers for RT-qPCR.

### Western blot analysis

Approximately 1 mg tissue is homogenized in 500 μL RIPA buffer (final concentrations: 50 mM Tris, 150 mM NaCl, 1% Triton-X100, 0.1% SDS, 0.5% sodium deoxycholate) with complete protease and phosphatase inhibitors (Roche), and samples were then clarified by centrifugation at 10,000g for 10 minutes. Total protein concentration was determined by BCA protein estimation (Thermo Scientific; Cat: 23225), and 30 μg of total protein was loaded onto 4–12% NuPAGE Bis-tris gels in MES buffer (Invitrogen; Cat: NP0323). After electrophoresis, proteins were transferred to 0.45 μm PVDF (Thermo Scientific; Cat: 88518). Membranes were blocked with 2% BSA-PBS and then incubated with Actin (Santa Cruz; SC-69879; 1:2000 dilution) and TBCK (Santa Cruz; SC-81865; 1:500 dilution) primary antibodies dissolved in 2% BSA-PBS overnight at 4 °C, followed by Li-Cor IRDye 800CW antibody (Li-Cor; 926-32210; 1:10000 dilution) for one hour at room temperature. Blots were developed with the Li-Cor Odyssey system.

### Micro-CT analysis

Tbck^+/+^, Tbck^-/-^ and Tbck^+/-^ mice were collected at E15, P0, P15, and adult (≥P60 for Tbck^+/+^ and Tbck^+/-^ mice as Tbck^-/-^ mice do not survive to adulthood) timepoints. After euthanasia with CO_2_, cervical dislocation, and decapitation (postnatal only), heads were stored 4% paraformaldehyde for 48 hours, then transitioned to 70% ethanol at 4°C until µCT images were obtained. To enhance visualization, embryos were stained according to published protocol with Phosphotungstic Acid (Lesciotto et al., 2020). A Scanco uCT35 (SCANCO Medical AG, Switzerland, Penn Center for Musculoskeletal Disorders MictoCT Imaging Core, Perelman School of Medicine, University of Pennsylvania) at a resolution of at least 15µm (higher resolution was employed for embryonic specimen). Skulls and embryos were reconstructed with Scanco software and analysis of images was performed with 3DSlicer (V5.2.2, Slicer.org) (Fedorov et al., 2012). Threshold settings were optimized to visualize only bone volume (postnatal), or only tissue volume (embryo). Virtual endocast of the skull (postnatal) and ventricles (embryo) were created to compare volumes using 3DSlicer Software and Wrap Solidify extension.

### Histology

After µCT analysis, P15 brains were carefully dissected from hard tissue and isolated brain tissue and whole embryos were processed for paraffin-based histology (CHOP Pathology Core). Prior to processing, P0 skulls were demineralized using 0.25 M EDTA at pH 7.4 for 12 days with three changes to ensure decalcification. Using a rotary microtome, 7µm coronal sections were cut and mounted on Superfrost-Plus slides (Thermofisher Scientific). Basic morphological staining proceeded according to previously published protocols (Behringer et al., 2013; Lesciotto et al., 2020; Motch Perrine et al., 2021; Pitirri et al., 2020). For immunohistochemistry, slides were subjected to epitope retrieval using a vegetable steamer and sodium citrate puffer pH 6.0 before blocking with 5% Goat serum in PBST. Primary antibodies were incubated overnight at 4°C **(Table 3)**. Secondary antibodies were added for 2 hours at room temperature, then counterstained with DAPI for 7 minutes before dehydration, clearing and mounting. Images were gathered using the Keyence BZ-X800 microscope. 20x images were stitched together using the Keyence BZ-X Series Analysis software. ImageJ (1.54g) was used to quantify immunofluorescent signals. Images were separated by channel, a standard ROI was added in the same anatomical region for all images, a standard threshold was used to remove background and staining artifact, and the number of cells positive for each target were counted using analyze particles (Sangree et al., 2025).

**Table 3:** Antibodies used for Immunohistochemical analysis

| Target | Manufacturer (Catalogue #) | Concentration |
| --- | --- | --- |
| GFAP | Antibodies Inc (75-240) | 1:250 |
| MAP2 | Proteintech (17490-1-AP) | 1:1250 |
| Calbindin | Antibodies Inc (75-488) | 1:250 |
| SOX2 | AbCam (Ab97959) | 1:200 |
| Iba1 | Invitrogen (PA5-27436) | 1:400 |
| NeuN | Sigma (MAB377) | 1:50 |
| Anti Mouse 488 | Invitrogen (A-21121) | 1:1000 |
| Anti Rabbit 647 | Invitrogen (A-21245) | 1:1000 |

### RNA-sequencing

Whole brain tissue was harvested from P0 mice with representation of male and female animals, in compliance with the Institutional Animal Care & Use Committee at the Children’s Hospital of Philadelphia. Tissue was snap frozen in liquid nitrogen and stored in –80C until RNA extraction. Total RNA was extracted from up to 20 mg of the snap-frozen tissue using the Promega simplyRNA Tissue Kit (AS1340) on the Maxwell® RSC Instrument (AS4500), following the manufacturer’s protocol, including on-cartridge DNase I treatment. RNA quantity and quality were assessed using a NanoDrop 8000 Spectrophotometer and the Agilent 4200 Bioanalyzer (TapeStation 4200 for RNA QC).

For library preparation, poly(A)-selected RNA libraries were generated using the Illumina Stranded mRNA Prep kit. A total of 300 ng of input RNA was used, with 11 cycles of PCR amplification. Libraries were sequenced on an Illumina NovaSeq 6000 system using an S1 flow cell with a 200-cycle configuration (paired-end 101 × 101 reads with dual 10 bp index reads). Library quality control was performed using the Revvity Gx HT Analyzer.

Raw sequencing read quality was assessed using FastQC and summarized with MultiQC. All samples exhibited high per-base sequence quality (median Phred scores >30), minimal adapter contamination, and GC content consistent with expectations, with no significant overrepresented sequences. Samples had a minimum of 13.7 million reads with a median of 15.4 million reads per sample. Reads were pseudoaligned using Kallisto, and downstream analyses were performed in R (version 4.4.0) using DESeq2 (version 1.44.0) for differential expression and ggplot2 (version 3.5.2) for visualization.

### STRING database protein functional enrichment analysis

Protein–protein interaction (PPI) analysis was performed using the STRING database (Search Tool for the Retrieval of Interacting Genes/Proteins, v12.0) (Szklarczyk et al., 2022). The list of differential expressed genes of interest was input, and interactions were retrieved using a minimum required interaction score of 0.4 (medium confidence) unless otherwise specified. Both physical and functional associations were included, derived from experimental data, curated databases, co-expression, and text mining.

### Statistics

Statistical analysis compared between genotypes (*Tbck^+/+^, Tbck^-/-^, Tbck^+/-^)* were made using Statskingdom (statskingdom.com) and R Studio software. Where assumptions for ANOVA were violated, non-parametric Kruskal Wallis Tests were employed. Where needed, assessments for outlier datapoints (due to histological artifact) were performed using Graphpad (Graphpad.com).

## Acknowledgements

We acknowledge the tremendous contributions of TBCK Warriors and their families without whom this would not be possible. Support from the TBCK Foundation (https://www.tbckfoundation.org/) and the TBCK Foundation of Puerto Rico was made possible by TBCK Warrior families and their direct contributions. We thank the Center for Applied Genomics (CAG) at the Children’s Hospital of Philadelphia (CHOP) for their support, including the CAG Sequencing Core who performed the sequencing analyzed here. Finally, we acknowledge the CHOP animal core and the University of Pennsylvania Neurobehavior Testing Core for their support of the animals in this investigation.

## Data and Resource Availability

All relevant data and details of resources can be found within the article and its supplementary information

## Author Contributions

All authors contributed to this work. Conceptualization: AJMP, ELD, DELC, EJKB

Methodology: AJMP, ELD, DELC, ADR, EMG, RA, SMS, KTW, KAK, BC, WTO, EJKB

Validation: AJMP, ELD, SMS, ADR, KTW, KAK, RA, XMW, BC, AB, WTO, DN

Formal analysis: AJMP, ELD, EMG, KJC, DELC

Investigation: AJMP, ELD, DELC, EMG, RA, SMS, KTW, KAK, BC, WTO, EJKB

Resources: WTO, EJKB Writing: ELD, EEL, AJMP, EMG

Editing: ELD, EEL, AJMP, EMG, DELC, SMS, XMW, ADR, BC, WTO

Visualization: AJMP, EMG, ELD, KJC, EEL, RA

Supervision: EJKB

Administration: EJKB

Funding: AJMP, ELD, DELC, EEL, KAK, EJKB

